# Crosstalk between enterocytes and innate lymphoid cell drives early IFN-γ-mediated control of *Cryptosporidium*

**DOI:** 10.1101/2021.03.13.435244

**Authors:** Jodi Gullicksrud, Adam Sateriale, Julie Englies, Alexis Gibson, Sebastian Shaw, Zachary Hutchins, Lindsay Martin, David Christian, Gregory A. Taylor, Masahiro Yamamoto, Daniel P. Beiting, Boris Striepen, Christopher A. Hunter

## Abstract

The intestinal parasite, *Cryptosporidium*, is a major contributor to global child mortality and causes opportunistic infection in immune deficient individuals. Innate resistance to *Cryptosporidium*, which specifically invades enterocytes, is dependent on the production of IFN-γ, yet whether enterocytes contribute to parasite control is poorly understood. In this study, utilizing the natural mouse pathogen, *Cryptosporidium tyzzeri*, we show that epithelial-derived IL-18 synergized with IL-12 to stimulate innate lymphoid cell (ILC) production of IFN-γ. This innate IFN-γ was required for early parasite control. Loss of STAT1 in enterocytes, but not dendritic cells or macrophages, antagonized early parasite control. Transcriptional profiling of enterocytes from infected mice identified an IFN-γ signature and enrichment of anti-microbial effectors like IDO, GBP and IRG. Deletion experiments identified a role for Irgm1/m3 in parasite control. Thus, enterocytes promote ILC production of IFN-γ that acts on enterocytes to restrict the growth of *C. tyzzeri*.

## INTRODUCTION

The intestinal epithelium is an important site for nutrient uptake and a barrier to micro-organisms. However, this barrier can be disrupted by a diverse group of viruses, bacteria and parasites that infect the gastrointestinal tract. The enteric diseases caused by these pathogens are of public health importance, accounting for 8-10% of deaths in children worldwide (Collaborators, 2016). While many of these pathogens will disseminate from the gut, a subset is restricted to the epithelial layer, where their interactions with enterocytes are likely key determinants of disease outcome. The epithelium is composed of enterocytes, a large population of columnar epithelial cells necessary for nutrient uptake, as well as subsets of specialized epithelial cells including Paneth, goblet and tuft cells that have roles in mucosal homeostasis and immune defense (Adolph et al., 2013; Birchenough et al., 2015; Cliffe et al., 2007; Gerbe et al., 2016; Nadjsombati et al., 2018; Nusse et al., 2018; Peterson and Artis, 2014; Schneider et al., 2018; von Moltke et al., 2016).

How enterocytes participate in resistance to different types of infection is a fundamental question that is particularly relevant to epithelial-restricted pathogens such as rotavirus, norovirus, astrovirus, *Shigella, Cyclospora* and *Cryptosporidium*. The apicomplexan parasite, *Cryptosporidium*, is a leading cause of severe diarrhea and death in infants (Checkley et al., 2015; Khalil et al., 2018; Kotloff et al., 2013; Platts-Mills et al., 2015) and a common opportunistic infection in individuals with primary and acquired immune deficiencies (Gomez Morales et al., 2004; Levy et al., 1997). Currently, there are no fully effective drugs or vaccines to treat or prevent cryptosporidiosis. Based on clinical experience and animal models, effective control and clearance of *Cryptosporidium* is dependent on cell-mediated immunity and the production of IFN-γ (Hayward et al., 2000; Leav et al., 2005; Tessema et al., 2009; White et al., 2000). *Cryptosporidium* invasive stages infect the apical surface of enterocytes, where they transform into the replicative trophozoite and occupy a unique intracellular but extracytoplasmic niche (Guerin and Striepen, 2020). The restriction of this parasite to the intestinal epithelium provides a model to understand how the immune system senses enteric pathogens and an opportunity to identify enterocyte-driven pathways that promote the control of intracellular pathogens.

Because *Cryptosporidium parvum*, the species most widely used as an experimental model, does not robustly infect adult immune competent mice, most studies have used immune deficient mice to help define important pathways for resistance to this organism. This approach has provided evidence that dendritic cell (DC) derived IL-12 promotes NK cell production of IFN-γ required for innate restriction of this infection (Barakat et al., 2009a; Bedi et al., 2014; Ehigiator et al., 2007; Potiron et al., 2019; Rohlman et al., 1993). IL-18 is another cytokine that promotes innate resistance to *C. parvum*. However, it is unclear if this is due to its ability to enhance ILC production of IFN-γ or whether IL-18 directly activates epithelial cells to limit parasite growth (Bedi et al., 2015; Choudhry et al., 2012; McDonald et al., 2006; McNair et al., 2018). Likewise, while IFN-γ is important for parasite control, multiple cell types in the gut can respond to this cytokine and it is unclear whether the ability of IFN-γ to activate enterocytes, dendritic cells or macrophages *in vivo* contributes to resistance to *Cryptosporidium*.

The recent description of the murine pathogen, *C. tyzzeri*, which is closely related to the species that infect humans, provides a natural experimental system to dissect the events required for protective immunity in an immunocompetent setting (Sateriale et al., 2019). In this study, infection with *C. tyzzeri* revealed that intestinal type-1 innate lymphoid cells (ILC1s) are a rapid source of IFN-γ that limits parasite growth. This protective response was dependent on the production of IL-12 and epithelial-derived IL-18. Lineagespecific deletion of STAT1, a critical transcription factor downstream of IFN-γ signaling, demonstrated that STAT1 was uniquely required in enterocytes—but not DCs or macrophages—to restrict parasite growth. Transcriptional profiling of enterocytes from infected mice highlighted an IFN-γ signature, and subsequent deletion experiments identified immune-related GTPase-m1 and −m3 (Irgm1/3) as downstream effectors of the IFN-γ-mediated response. Together, these studies establish that enterocytes have a central role as a source of IL-18 required to stimulate local ILC responses and are key mediators of IFN-γ-mediated protection to an important enteric pathogen.

## MATERIALS & METHODS

### Mice

C57BL/6 (stock no: B6NTac), Rag2−/− (stock no: RAGN12), and Rag2−/−Il2rc−/− (stock no: 4111) were purchased from Taconic. C57BL/6 (stock no: 000664), Rag2−/− (stock no: 008449), Ifng−/− (stock no: 002287), IL-12p40 KO (stock no: 002693), Stat1−/− (stock no: 012606), II18−/− (stock no: 004130), Ido1−/− (stock no:005867), Vil1-Cre (stock no:021504), LysM-Cre (stock no: 004781), and Cd11c-Cre (stock no. 008068) were purchased from Jackson Laboratory and maintained in-house. STAT1flox mice were generated as previously described (Klover et al., 2010) and maintained in house. Ifng/Thy1.1 BAC-In mice were provided by Dr. Phillip Scott but originated in the laboratory of Dr. Casey Weaver (Harrington et al., 2008; Hatton et al., 2006). Il18-flox mice were provided by Dr. Jorge Henoa-Mejia. Vil1-Cre^ERT2^ were provided by Dr. Lou Ghanem at Children’s Hospital of Philadelphia and Dr. David Artis at Cornell University. In-house breeding was performed to obtain all Cre-lox combinations. Unless otherwise noted, mice used in this study were males or females ranging from 7-11 weeks. We did not observe a difference in infection burden between male and female mice. All mice were age matched within individual experiments. All protocols for animal care were approved by the Institutional Animal Care and Use Committee of the University of Pennsylvania (protocol #805405 and #806292).

### Parasites and infection

Transgenic *C. tyzzeri* expressing nanoluciferase and mCherry (Sateriale et al., 2021; Sateriale et al., 2019) are propagated by orally infecting *Ifng*^−/−^ mice. Oocysts are purified from fecal collections of infected mice using sucrose flotation followed by a cesium chloride gradient, as previously described (Sateriale et al., 2019). Mice were infected with 5×10^4^-1×10^5^ oocysts by oral gavage. To measure parasite burden in intestinal tissue, 5mm biopsy punches were taken along murine small intestines and suspended in 0.5mL lysis buffer (50mM tris HCl (pH 7.6), 2mM DTT, 2mM EDTA, 10% glycerol, 1% TritonX in ddH_2_O). To quantify fecal oocyst shedding, 20mg fecal material was suspended in 1mL lysis buffer. Samples were shaken with glass beads for 5min, then combined in a 1:1 ratio with Nano-Glo® Luciferase solution (Promega, Ref N1150). A Promega GloMax plate reader was used to measure luminescence.

### Histology

For histological analysis of the small intestine, tissue from the distal third of the small intestine was flushed with 10% neutral buffered formalin (Sigma, St Louis, MO, USA), then ‘swiss-rolled’ and fixed overnight. Fixed samples were paraffin-embedded, sectioned, and stained with hematoxylin and eosin for detailed histologic evaluation. Slides were evaluated by a board-certified veterinary pathologist in a blinded fashion for quantitative measurements of number of parasites, villus/crypt architectural features and inflammatory infiltrates, and semi-quantitative scores for villus epithelium lesions as previously described (Sateriale et al., 2019).

### Cytokine neutralization and measurement

To neutralize IFN-γ, 1mg anti-IFN-γ (XMG1.2, BioXcell Cat #: BE0055) was given intraperitoneally (i.p.) 1 day prior and 2 days post infection with *C. tyzzeri*. To also neutralize IL-12 or IL-18, 2mg anti-IL-12p40 (C17.8 BioXcell Cat #: BE0051) or 1mg anti-IL-18 (YIGIF74-1G7, BioXcell Cat #: BE0237) were given i.p. on days −4, −1, and 1, while anti-IFN-γ was given on day −2 and day 2. Intestinal IFN-γ levels were assessed from 5mm biopsy punches that were incubated in complete RPMI at 37°C for 24 hours. Clear, flat-bottom 96-well plates (Immunulon 4 HBX) were coated with 0.25μg/mL anti-IFN-γ (AN-18, Invitrogen Ref #:14-7313-85) at 4°C overnight. Samples were added and IFN-γ left to bind at 37°C for 2 hours. 0.25μg/mL biotinylated anti-IFN-γ (R4-6A2, eBioscience Ref #: 13-7312-85) in PBS with 2.5% FBS and 0.05% Tween was added for 1 hour at room temperature, followed by peroxidase-labeled streptavidin for 30 minutes. Finally, KPL ABTS® peroxidase substrate (SeraCare Cat #: 5120-0041) was applied for detection.

### Flow cytometry and cell sorting

Single-cell suspensions were prepared from intestinal sections by shaking diced tissue at 37C for 25 minutes in Hank’s Balanced Salt Solution with 5 mM EDTA and 1 mM DTT. Cell pellets were then passed through 70 mm and 40 mm filters. Cells were surface stained using the following fluorochrome-conjugated Abs: anti-EpCAM (G8.8), anti-CD45.2 (104), anti-CD19 (MB19-1), anti-Thy1.1 (HIS51), anti-CD27 (LG.7F9), anti-RORγt (B2D) and anti-γδ TCR (eBioGL3) from eBioscience; anti-NK1.1 (PK136), anti-NKp46 (29A1.4), anti-CD3 (17A2), anti-CD127 (SB/199), anti-CD49a (HMα1), anti-T-bet (4B10) and anti-CD8ß (YTS156.7.7) from BioLegend; anti-Eomes (Dan11mag) and Live/Dead Aqua from Invitrogen; and anti-CD49b (DX5) from BD Biosciences. Data were collected on a LSRFortessa (BD Biosciences) and analyzed with FlowJo v10 software (TreeStar). Cell sorting was performed on a BD FACS Jazz (BD Biosciences).

### RNA-seq and gene enrichment analysis

RNAseq reads were pseudo-aligned to the Ensembl *Mus musculus* reference transcriptome v79 using Kallisto v0.44.0 (Bray et al., 2016). In R, transcripts were collapsed to genes using Bioconductor tximport (Robinson et al., 2010), and differentially expressed genes were identified using Limma-Voom (Law et al., 2014; Ritchie et al., 2015). Gene set enrichment analysis was performed using the GSEA software and the annotated gene sets of the Molecular Signatures Database (MSigDB) (Mootha et al., 2003; Subramanian et al., 2005). From the GSEA output, enrichment maps were generated to provide a visual representation of gene set overlap using Cytoscape v3.8.2 (Shannon et al., 2003). Data and analyses have been deposited to the GEO repository (GSE168680).

### Statistics

Statistical significance was calculated using the unpaired Student’s t-test for comparing 2 groups, or ANOVA followed by multiple comparisons for comparing groups of 3 or more. Analyses were performed using GraphPad Prism v.9.

## RESULTS

### Localized innate IFN-γ provides early constraint of *C. tyzzeri* infection

To assess the relationship between parasite burden and IFN-γ production, WT mice treated with an isotype control antibody (IgG) or anti-IFN-γ (α-IFN-γ) were infected with a *C. tyzzeri* strain that expressed nanoluciferase (Sateriale et al., 2021). At 4 days post-infection (dpi), paired biopsies were taken along the entirety of the intestine and used to quantify nanoluciferase activity or placed in culture to assess levels of secreted IFN-γ. In control mice, parasite replication was restricted to the distal small intestine but levels of IFN-γ in ileal supernatants were not elevated above uninfected controls (Figure 1A). However, in mice treated with anti-IFN-γ, there was an 80-fold increase in parasite burden and *ex vivo* IFN-γ was readily detected and correlated with areas of the gut with the highest parasite burdens (Figure 1A, filled circles). Thus, the early production of IFN-γ provides a mechanism of resistance to *C. tyzzeri*, but the ability to detect the infection-induced production of IFN-γ is dependent on the presence of a high tissue parasite burden.

**Figure 1.**
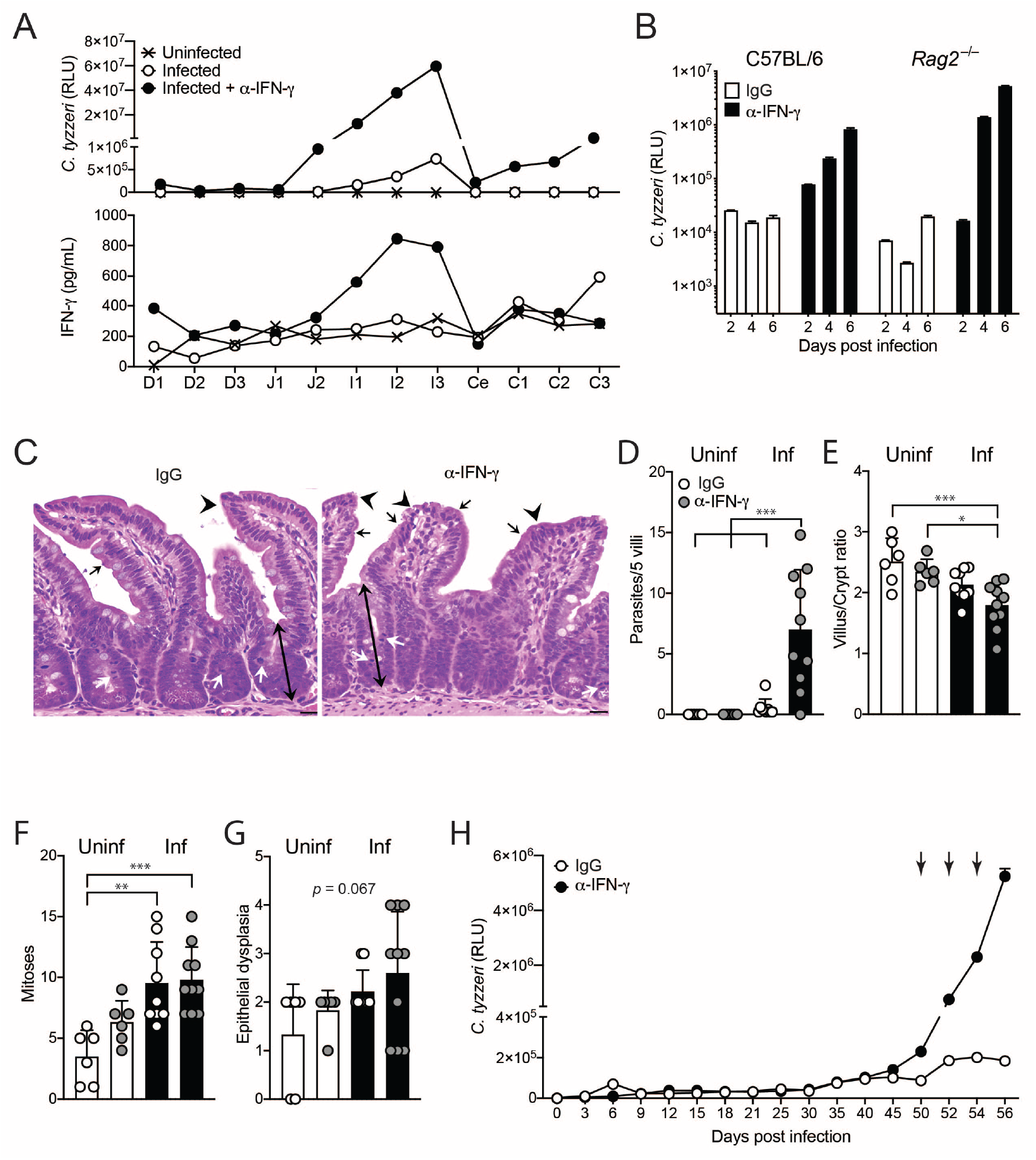
Localized innate IFN-γ provides early protection from *C. tyzzeri* and pathology. WT mice were infected with 10^5^ *C. tyzzeri* oocysts and treated with either α-IFN-γ or isotype control. (A) At 4dpi, two adjacent 5mm biopsies were taken along the length of the small and large intestines (D=duodenum, J=jejunum, I=ileum, Ce=cecum, C=colon). One biopsy was evaluated for nanoluciferase (top), the other was incubated at 37°C for 24h and IFN-γ was measured from the supernatant. Graph depicts 1 representative mouse from n=3 for each group. (B) WT C57BL/6 and *Rag2*^−/−^ mice were treated with isotype or α-IFN-γ and nanoluciferase was used to measure fecal oocyst shedding, n=4 per group. Similar results were observed in 2 additional experiments. (C) Representative mucosa and (D-G) cumulative histology scoring from *Rag2*^−/−^ mice 5dpi that shows a marked increase in cryptosporidia organisms (black arrows) infecting the villus enterocytes in treated mice versus controls with a progressive increase in epithelial dysplasia (arrowheads), reduced villus:crypt ratios with increased crypt depth (double-headed arrows) and crypt mitoses (white arrows); scale bars = 20μm. (H) *Rag2*^−/−^ mice were infected with 10^5^ *C. tyzzeri* oocysts and treated with α-IFN-γ at 50, 53, and 56dpi. Oocysts shedding was monitored throughout by nanoluciferase. n=4, representative from 3 experimental replicates. Bars denote mean ± SD. ANOVA followed by multiple comparisons were performed on cumulative pathology scores, \*\**p* ≤0.01, \*\*\**p* ≤ 0.001.

To assess the contribution of innate and adaptive sources of IFN-γ on early resistance to *C. tyzzeri*, WT and *Rag2*^−/−^ mice (lacking T and B cells) were treated with α-IFN-γ or an isotype control prior to infection. No gross histological differences were detected between mouse strains at steady state (Supplemental Figure 1A). In both WT and *Rag2*^−/−^ mice, IFN-γ neutralization resulted in increased oocyst shedding by 4dpi, which was exacerbated at 6dpi (43-fold in WT and 263-fold in *Rag2*^−/−^, Figure 1B), and correlated with histological analysis that showed enhanced parasite burden in *Rag2*^−/−^ mice treated with α-IFN-γ (Figure 1 C-D). In addition, there were significant infection-induced local changes in mucosal architecture and enterocyte morphology by 5dpi, including reduced average villus:crypt ratio, increased crypt mitoses and a trend toward increased epithelial dysplasia (Figure 1E-G). However, quantification of these changes across multiple sections did not yield statistically significant changes between infected mice and those treated with α-IFN-γ. However, when observing ileal pathology at a later timepoint, α-IFN-γ treatment resulted in more severe pathological changes (12dpi Supplemental Figure 1B). The pathology corresponds with the sustained oocyst shedding observed after the peak parasite burden at 6dpi. Interestingly, treatment of chronically infected *Rag2*^−/−^ mice with α-IFN-γ at 50dpi led to a marked recrudescence in parasite burden (Figure 1H). These data identify an innate mechanism of IFN-γ-mediated resistance to *C. tyzzeri* that operates during the acute and chronic phases of infection, but which is insufficient for parasite clearance.

### Innate lymphoid cells are required for control of *C. tyzzeri*

There are several ILC populations capable of IFN-γ production that include: NK cells, ILC1s and ILC3s. Early studies with *C. parvum* concluded that NK cells were a major source of IFN-γ, but the use of α-asialo-GM1 (which depletes NK cells but not other ILC populations) indicated that there may be other innate sources of IFN-γ (Rohlman et al., 1993; Ungar et al., 1991). Indeed, when *Rag2*^−/−^ mice infected with *C. tyzzeri* were treated with α-asialo-GM1, there was efficient depletion of splenic NK cells but the intestinal ILCs (CD45.2^+^ NK1.1^+^NKp46^+^) and levels of secreted IFN-γ were only reduced by approximately half (Supplemental Figure 1C-E). Therefore, to assess the role of ILCs in resistance to *C. tyzzeri, Rag2*^−/−^Il2rg^−/−^ mice, which lack all ILCs in addition to T and B cells, were compared to WT and *Rag2*^−/−^ mice. After infection with *C. tyzzeri*, oocyst shedding declined in WT mice by 9dpi, and resolved by 18dpi (Figure 2A). Parasite levels in *Rag2*^−/−^ mice were not elevated compared to WT mice at early time points (3-9dpi), but the infection failed to resolve in the absence of adaptive immune cells. In contrast, *Rag2*^−/−^/Il2rg^−/−^ mice demonstrated little evidence of parasite control at any time examined, with approximately 250-fold higher nanoluciferase readings by 6dpi, compared to *Rag2*^−/−^ mice, which were sustained for the duration of the experiment (Figure 2A). In uninfected mice, there were no gross histological differences in the ileal tissues among the three strains (Supplemental Figure 1A). At 5dpi, consistent with the data in Fig 1B, WT and *Rag2*^−/−^ mice had similar numbers of parasites and levels of infection-induced changes to the epithelium (Figure 2D-G, white and gray bars). In contrast, *Rag2*^−/−^/*Il2rg*^−/−^ mice exhibited dramatically increased parasite burden (Figure 2B-C). The absence of ILCs also resulted in increased mitoses and crypt branching, as well as more severe villus pathology, indicated by increased epithelial dysplasia and attenuation (Fig 2D-G, black bars). Thus, in the absence of adaptive immunity, ILCs are required for both early and long-term control of *C. tyzzeri* and the lack of these cells results in unrestricted parasite replication and severe intestinal pathology.

**Figure 2.**
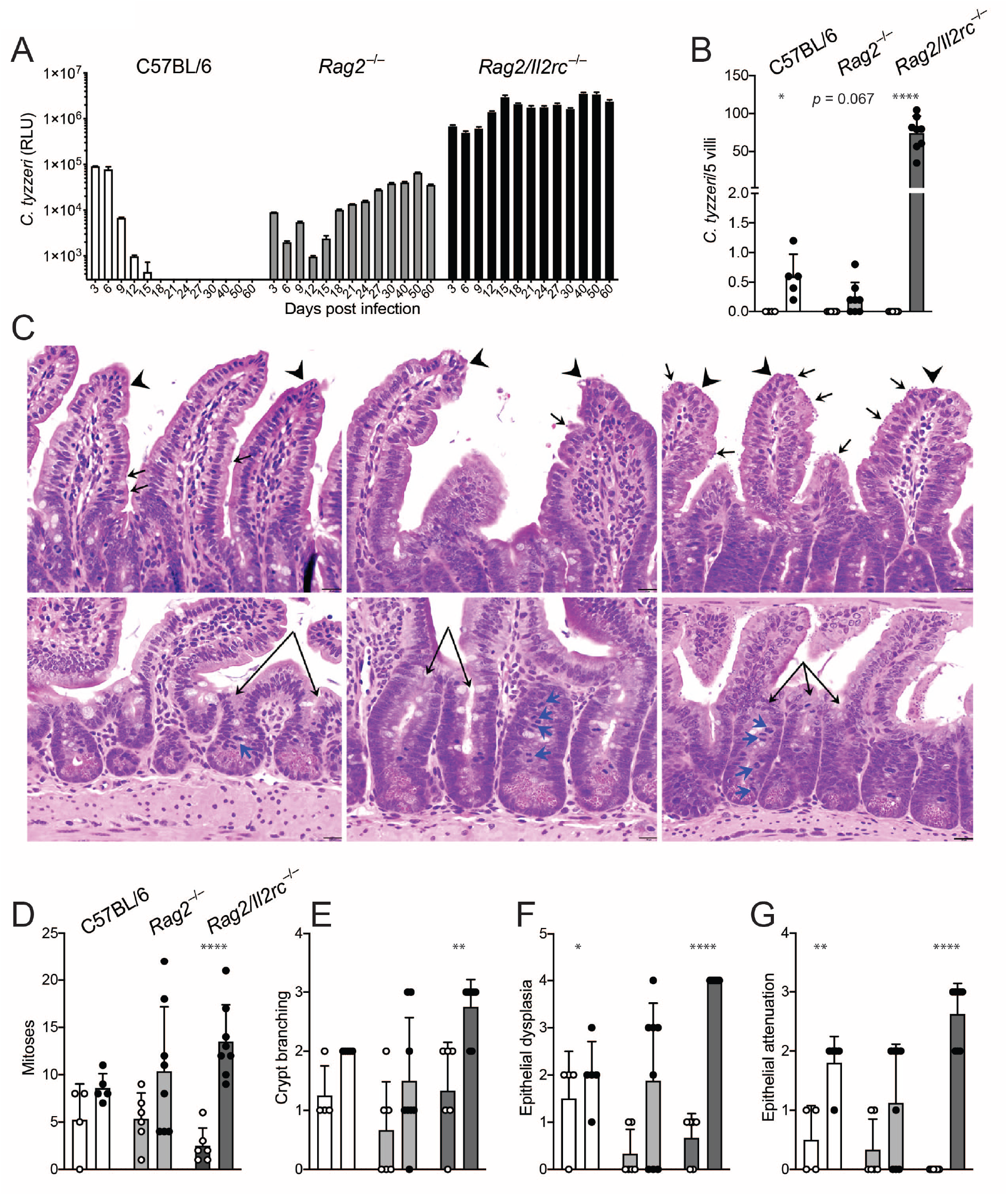
Innate lymphoid cells are critical components of protection against *C. tyzzeri*. C57BL/6 WT, *Rag2*^−/−^ and *Rag2*^−/−^*Il2rg*^−/−^ mice were infected and used for histological analysis or oocyst shedding. (A) Kinetics of fecal oocyst shedding; n=3-5, representative of 3 replicates comparing *Rag2*^−/−^ and *Rag2*^−/−^ *Il2rg*^−/−^. (B-G) H&E staining of representative villi (upper panels) and crypts (lower panels), and cumulative histology scoring from WT, *Rag2*^−/−^ and *Rag2*^−/−^*Il2rg*^−/−^ mice at 5dpi that show a marked increase in *Cryptosporidium* organisms (black arrows) infecting the villus enterocytes in *Rag2*^−/−^Il2rg^−/−^ mice versus WT or *Rag2*^−/−^ mice as well as progressive villus epithelial dysplasia with attenuation (arrowheads), increases in crypt branching (double arrows) and crypt mitoses (blue arrows); scale bars = 20μm; n = 4-8 where each symbol denotes 1 mouse cumulative of 3 experiments. Bars indicate mean ± SD. n = 4-8 where each symbol denotes 1 mouse, cumulative of 3 experiments. Bars indicate mean ± SD for uninfected (open circles) or 5dpi (filled circles). For histology scoring, t-tests were used to compare uninfected and infected within each mouse strain, *\*p* ≤ 0.05, \*\**p* ≤ 0.01, \*\*\*\**p* ≤ 0.0001.

### ILC1s are a major source of innate IFN-γ during *C. tyzzeri* infection

To determine the innate cellular source(s) of IFN-γ in the intestinal epithelium, mice in which the gene for the surface-expressed protein, Thy1.1, is under the transcriptional control of the *Ifng* promotor (Harrington et al., 2008) were infected with mCherry-expressing *C. tyzzeri*. This approach allowed the simultaneous quantification of parasite-infected (mCherry^+^) enterocytes (EpCAM^+^CD45^-^) and innate lymphoid cells (CD45.2^+^CD3^-^CD19^-^NK1.1^+^NKp46^+^) producing IFN-γ (Thy1.1^+^). Based on the data in Fig 1A, groups of mice were also treated with α-IFN-γ to increase the ability to detect cells that produce IFN-γ. In uninfected mice, enterocytes lack mCherry, and a low basal percentage of ILCs express surface Thy1.1 regardless of α-IFN-γ treatment (Figure 3A, C). At 4dpi, a detectable portion of IECs were infected (mCherry^+^), and there was a small increase in the frequency of Thy1.1^+^ ILCs. Treatment with α-IFN-γ resulted in a significant increase in parasite burden (Figure 3A-B) and a 3-5-fold increase in the proportion of Thy1.1^+^ ILCs (Figure 3C-D). This increased production of IFN-γ was specific to the intestinal epithelium, as few cells expressed Thy1.1 in Peyer’s patches or mesenteric lymph nodes, even with IFN-γ neutralization (Figure 3E-F). Of note, in these immune competent reporter mice, there was also infection-induced expression of Thy1.1 by intestinal CD4^+^ T cells and, to a lesser extent, CD4^+^CD8α^+^ T cells (Supp Fig 2A).

**Figure 3.**
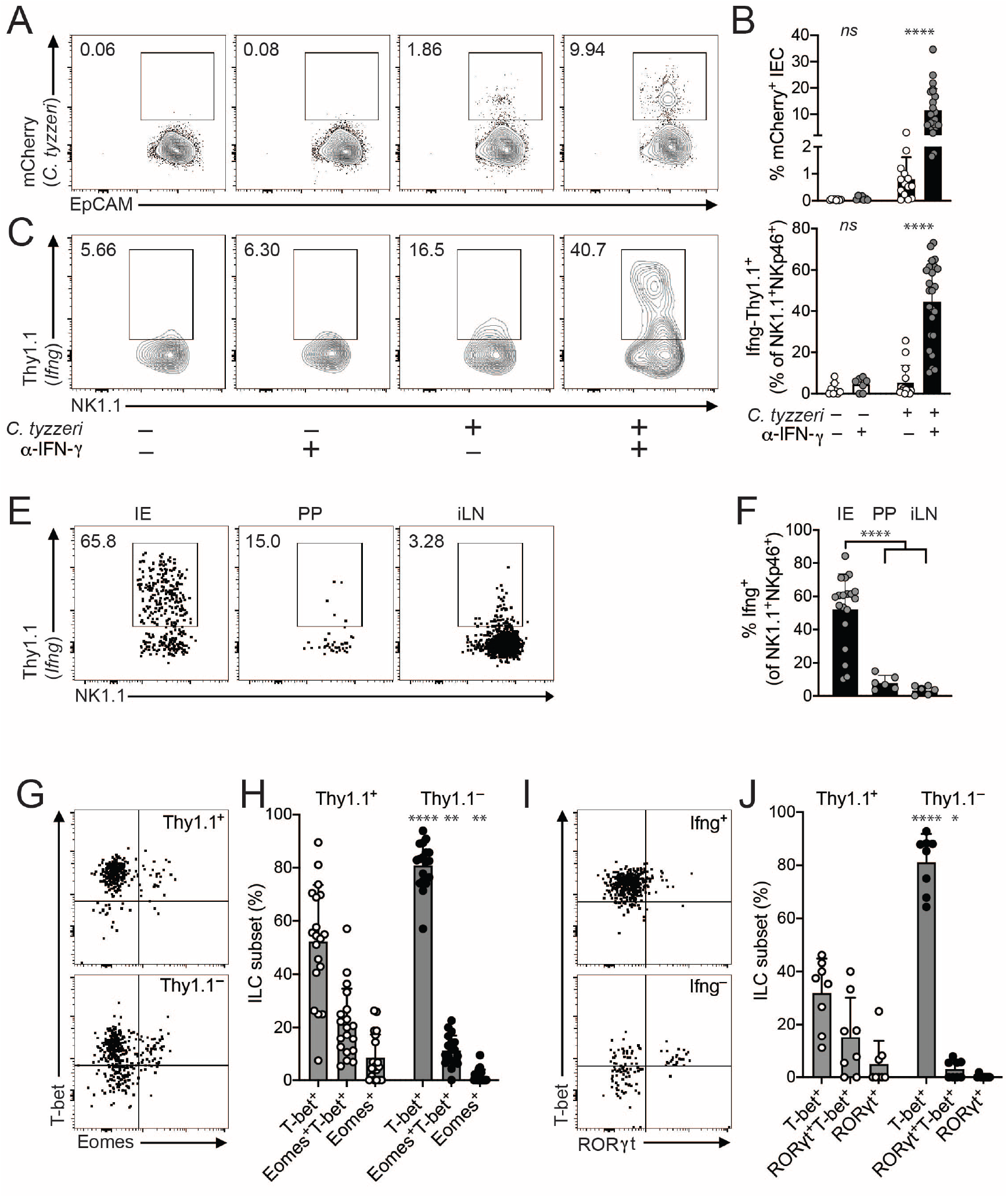
ILC1s are a critical source of innate IFN-γ. *Ifng*-Thy1.1 KI mice were infected with mCherry-expressing *C. tyzzeri* and cells from the ileal epithelium were assessed by flow cytometry at 4dpi. (A) Representative flow plots and (B) summary bar graph of *C. tyzzeri*-infected (mCherry^+^) intestinal epithelial cells, gated on Live CD45.2^-^EpCAM^+^ cells. (C) Representative flow plots and (D) summary bar graph of Ifng^+^ (Thy1.1^+^) innate lymphoid cells (NK1.1^+^NKp46^+^), n = 7-24 from 4 experiments. (E) Representative flow plots and (F) summary bar graph of frequency of Ifng+ (Thy1.1) cells from ILCs (seeing gating strategy from (C)) taken from intestinal epithelium (IE), Peyer’s patches (PP), or mesenteric lymph nodes (mesLN). mesLN and PP were pooled from 2-4 mice, representative of 3 experiments. (G) Representative flow plots and (H) summary bar graph of T-bet and Eomes expression on Ifng^+^ (Thy1.1^+^) or Ifng^-^ (Thy1.1^-^) ILCs. (I) Representative flow plots and (J) summary bar graph of T-bet and RORγt expression on Ifng^+^ (Thy1.1^+^) or Ifng^−/−^ (Thy1.1^−/−^) ILCs. All ILCs were gated on Live CD45.2^+^CD3^-^CD19^-^NKp46^+^NK1.1 ^+^ cells. Statistical significance between α-IFN-γ treated or untreated in B and D was determined by Student’s t-test. Statistical significance in F was determined by one-way ANOVA and multiple comparisions. Student’s t-tests were used to compare Thy1.1^+^ and Thy1.1^-^ in H and J. ns = not significant (*p* > 0.05), \**p* ≤ 0.05, \*\**p* ≤ 0.01, \*\*\*\**p* ≤ 0.0001.

To characterize the ILC subsets that produce IFN-γ, expression of the transcription factors Eomes, T-bet and RORγt was used to distinguish NK cells, ILC1s, and ILC3s, respectively. In uninfected reporter mice, the ILC populations in the gut were composed of 50-60% T-bet^+^ ILC1s, 30-40% Eomes^+^ NK cells and 5-10% were RORγt^+^ ILC3s (Supplemental Figure 2B). All three of these ILC subsets were also present at 4dpi (Figure 3G and 3I). Among ILCs not expressing IFN-γ (Thy1.1^-^), the proportions of NK cells, ILC1s and ILC3s were similar to those of uninfected mice (Figure 3G, I, bottom plots and Figure 3H, J open circles). However, in infected mice, the proportion of Tbet+ ILCs was significantly increased in cells expressing *Ifng* (Thy1.1^+^), where approximately 80% of the Thy1.1^+^ cells expressed T-bet and not Eomes or RORγt (Figure 3G-J, filled circles). In addition, while NK cells in the mesenteric lymph nodes expressed CD27 and CD49b, the Thy1.1^+^ cells in the intestinal epithelium showed low expression of these NK cell markers (Supplemental Figure 2C-D). These data indicate that within days of *C. tyzzeri* infection, ILC1s present in the intestine become activated and are a major source of IFN-γ.

### Epithelial-derived IL-18 synergizes with IL-12 to promote ILC production of IFN-γ

IL-12 and IL-18 have roles in innate resistance to *Cryptosporidium sp*. (Bedi et al., 2015; Ehigiator et al., 2007; Sateriale et al., 2021) and can synergize to stimulate ILC production of IFN-γ in other experimental systems (Fuchs et al., 2013; Hunter et al., 1997; Takeda et al., 1998). Therefore, to evaluate their contributions to early resistance to *C. tyzzeri*, the level of oocyst shedding in WT, *Ifng*^−/−^, *Il12b*^−/−^ (IL-12p40) and *Il18*^−/−^ mice were compared. As expected, infection was established in WT mice, and *Ifng*^−/−^ mice showed a rapid and marked increase in parasite burden. While the parasite levels in *Il12b*^−/−^ mice were comparable to *Ifng*^−/−^ mice, *Il18*^−/−^ mice showed a phenotype that was intermediate between WT and *Ifng*^−/−^ mice (Figure 4A). A recent study highlighted BATF3-dependent CD103^+^ DCs as a source of IL-12 during neonatal infection with *C. parvum* (Potiron et al., 2019), but the cellular source of IL-18 required to control *Cryptosporidium* was unclear. To determine the importance of the intestinal epithelium as a cellular source of IL-18 during cryptosporidiosis, oocyst shedding from mice bearing an epithelial lineagespecific deletion of IL-18 (*Villin-Cre x Il18^fl/fl^*, here referred to as *Il18*^DIEC^) was compared to WT and *Il18*^−/−^ mice. In these studies, the levels of infection in *Il18*^DIEC^ mice were comparable to whole-body *Il18*^−/−^ mice (Figure 4B), suggesting that the intestinal epithelium is a key source of the IL-18 that mediates early resistance to *C. tyzzeri*.

**Figure 4.**
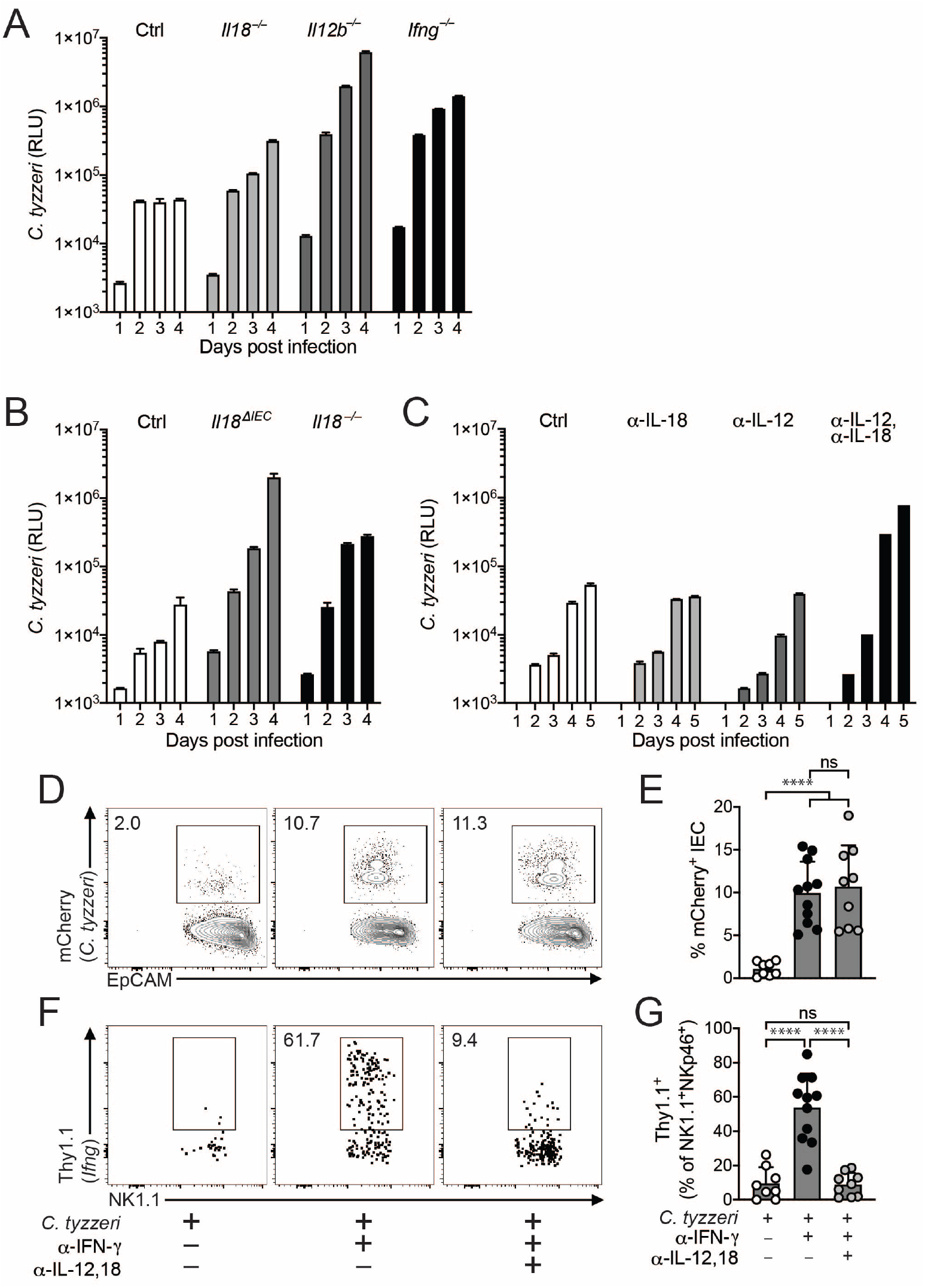
IL-12 and IL-18 stimulate IFN-γ production by ILCs. (A) WT C57BL/6 mice or mice lacking IL-18 (*Il18*^−/−^), IL-12p40 (*Il12b*^−/−^) or IFN-γ (*Ifng*^−/−^) were infected with *C. tyzzeri* and oocyst shedding in feces was monitored. Representative of 3 experimental replicates. (B) Control (Cre^-^ *Il18*^fl/fl^), epithelial-specific IL-18-deficient (Villin^Cre^ x *Il18^fl/fl^*; Il18^ΔIEC^) and *Il18*^−/−^ mice were infected with *C. tyzzeri* and oocyst shedding in feces was monitored. (C-G) *Rag2*^−/−^ mice (C) or *Ifng*-Thy1.1 KI mice (D-G) were treated with α-IL-18, α-IL-12p40, or both and infected with *C. tyzzeri*. (C) Oocyst shedding in feces was monitored. (D) Representative flow plots and (E) summary bar graph of *C. tyzzeri*-infected (mCherry+) IECs 4dpi. (F) Representative flow plots and (G) summary bar graph of Ifng^+^ (Thy1.1^+^) ILCs 4dpi (gating same as Figure 3). n = 8-11 from 3 experiments. Statistical significance was determined by one-way ANOVA and multiple comparisions; ns = not significant (*p* > 0.05) \*\*\*\**p* ≤ 0.0001.

The experiments that utilized mice with germline deletion of IL-18 or IL-12 identified an important role for these cytokines in early resistance to *C. tyzzeri*. However, analyses of naïve *Il12b*^−/−^ mice revealed a marked absence of ILC populations in the intestinal epithelium (Supp Fig 3A-C) and previous studies have reported that germline deletion of IL-18 impacts immune homeostasis in the gut (Harrison et al., 2015). In order to control for these confounding effects, C57BL/6 and *Rag2*^−/−^ mice were treated with antibodies against IL-12p40, IL-18, or both beginning 4 days prior to infection with *C. tyzzeri*. In WT and *Rag2*^−/−^ mice, neither treatment alone dramatically impacted susceptibility to *C. tyzzeri* but the simultaneous blockade of IL-12 and IL-18 led to a 14.5-fold increase in parasite burden (*Rag2*^−/−^: Figure 4C, BL/6: Supp Fig. 3D). Moreover, despite having similar infection burdens to α-IFN-γ treated mice (Figure 4D-E), *Ifng*-Thy1.1 reporter mice treated with α-IL-12p40 plus α-IL-18 showed nearly complete loss of Thy1.1^+^ ILC (Figure 4F-G). Collectively, these experiments demonstrate that epithelial-derived IL-18 synergizes with IL-12 to stimulate ILC production of IFN-γ required for early restriction of *C. tyzzeri* infection.

### Enterocytes are critical contributors to early IFN-γ-mediated resistance to *C. tyzzeri* infection

While IFN-γ is important in acute resistance to *C. tyzzeri*, it is unclear whether it acts directly on infected cells to restrict parasite growth, or indirectly via activation and maturation of macrophages and dendritic cells (Laurent and Lacroix-Lamande, 2017). The transcription factor STAT1 is the major mediator of IFN signaling. *Ifng*^−/−^ and *Stat1*^−/−^ mice had similarly enhanced susceptibility to *C. tyzzeri*, suggesting that IFN-γ is the main driver of STAT1 signaling required for resistance to this parasite (Supplemental Figure 4A). Therefore, lineage-specific Cre-lox mediated deletion was utilized to identify cell subsets in which STAT 1 signaling was critical for early control of *C. tyzzeri*. Loss of STAT 1 only in dendritic cells (*Cd11c-Cre* x *Stat1*^fl/fl^; *Stat1*^ΔDC^) or macrophages (*Lys2-Cre* x *Stat1*^fl/fl^; *Stat1*^ΔMΦ^) did not enhance parasite burden over Cre^-^ controls. In contrast, when STAT1 was deleted from enterocytes using a tamoxifen-inducible deletion (*Villin*-Cre^ERT2^ x *Stat1*^fl/fl^; *Stat1*^ΔIEC^), oocyst shedding was greatly increased (Figure 5A) and at a level that paralleled complete *Stat1*^−/−^ mice (Figure 5B). This was not due to reduced levels of IFN-γ in the intestine (Supp Fig. 4B). Thus, IFN-γ induced STAT1-mediated activity in intestinal epithelial cells provides a cell intrinsic mechanism for the control of *Cryptosporidium*.

**Figure 5.**
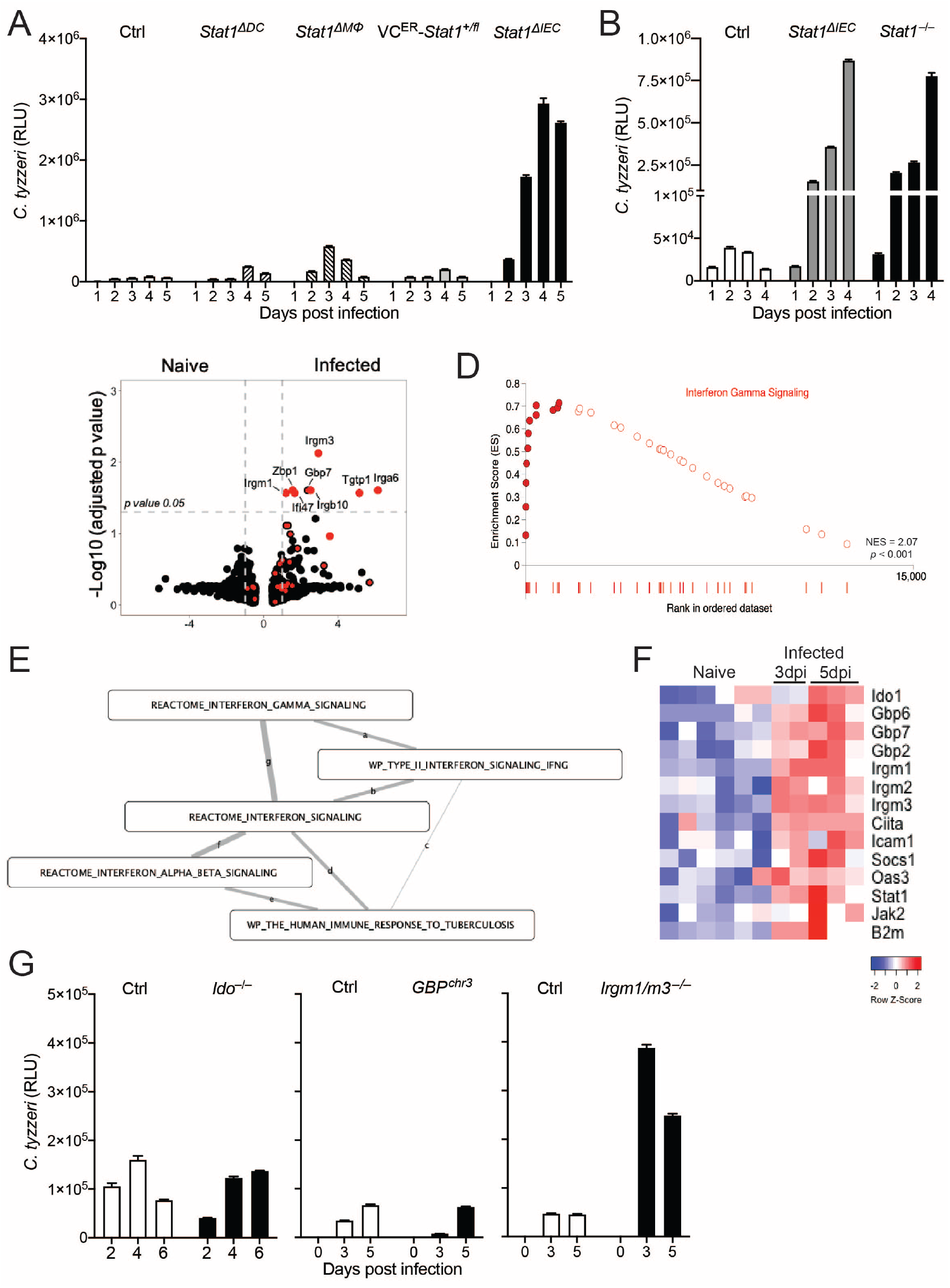
IFN-γ-mediated protection is dependent on enterocyte expression of STAT1. (A-B) Cre^-^Stat1^fl/fl^ (Control), Cd11c^Cre^Stat1^fl/fl^ (*Stat1DDC*) and Lyz2^Cre^Stat1^fl/fl^ (*Stat1*^ΔMΦ^), Villin^CreERT2^-Stat1^fl/fl^ (*Stat1*^ΔIEC^) and *Stat1*^−/−^ mice were infected with *C. tyzzeri* and oocyst shedding was monitored in feces. (C-F) IEC from naïve mice or mice infected with *C. tyzzeri* for 3 or 5 days were FACS sorted and used for RNA-sequencing analysis. (C) Volcano plot with IFN signature genes marked in red. DEGs that were significantly upregulated (*p* ≤ 0.05) in IECs from *C. tyzzeri*-infected mice are labeled. (D-E) Gene set enrichment analysis highlighting an IFN-γ signature (D), a cluster of IFN pathways (E). (F) Heatmap of pertinent IFN-stimulated genes. (G) WT C57BL/6, *Ido1*^−/−^, GBP^chr3^, and *Irgm1/m3*^−/−^ mice were infected with 5×10^4^ *C. tyzzeri* oocysts and fecal shedding of oocysts was monitored. Representative of 3 experimental replicates.

Because the mechanism that leads to the IFN-γ-mediated restriction of *Cryptosporidium* in enterocytes is unknown, transcriptional profiling of sort-purified enterocytes from uninfected or infected mice was performed. RNA-seq analysis of enterocytes identified a limited set of differentially expressed genes; that included several IFN-inducible GTPases: *Igtp* (encoding Irgm3), *Iigp1* (encoding Irga6), *Gm12250* (encoding Irgb10), *Gbp7, Tgtp1, Ifi47* (encoding Irg-47), and *Irgm1* (Figure 5C). Despite the limited number of differentially expressed genes, GSEA showed a strong enrichment of IFN-γ signaling in enterocytes from infected ileums (Figure 5D). Enrichment mapping software was used to define related gene functions and examine overlap between enriched gene sets. This analysis identified 3 main clusters: IFN signatures, mitochondrial signatures, and protein translation/ribosome signatures (Figure 5E and Supp Fig 4C-D). The mitochondrial and ribosome signatures have been associated with enterocyte stress (Moon, 2011; Rath et al., 2018) and likely reflect the increased mitotic index and epithelial dysplasia associated with this infection (see Figures 1 and 2). Furthermore, while gene sets encompassing both IFN-γ and type I interferons were enriched, the degree of overlap was greatest among the all-encompassing “interferon signaling” and the two interferon gamma gene sets, further indicating that IFN-γ signaling was the dominant response in infected IECs.

Given the dominant IFN-γ signature, the expression data sets were curated for changes in IFN-γ-induced genes associated with control of intracellular pathogens (Fig 5F). Genes strongly upregulated by *C. tyzzeri* infection included those encoding for ß2m and CIITA (Class II Major Histocompatibility Complex Transactivator) which affect MHC I and MHC II expression, respectively (Figure 5F). Also markedly upregulated were transcripts encoding for indolamine dioxygenase (IDO), several Guanylate Binding Proteins (GBPs) and Immunity Related GTPases (IRGs), all of which are known to be important in non-hematopoietic cells for IFN-γ to restrict the growth of *T. gondii*(Saeij and Frickel, 2017; Yamamoto et al., 2012). However, their roles in resistance to *Cryptosporidium* have not been established. To test the impact of these pathways on *C. tyzzeri*, the course of infection in *Ido1*^−/−^, GBP^chr3^ (lacking *Gbp2*, *Gbp3*, *Gbp5*, *Gbp7*, and *Gbp2ps*) and *Irgm1/m3*^−/−^ mice was compared with that in WT mice by fecal oocyst shedding. Because Irgm1-deficient mice have multiple defects in immune function that are mitigated by the loss of Irgm3, *Irgm1/m3*^−/−^ mice were used to study to role of Irgm1 in resistance to infections (Liu et al., 2013; Maric-Biresev et al., 2016). While *Ido1*^−/−^ and GBP^chr3^ mice showed no enhanced susceptibility to *C. tyzzeri, Irgm1/m3*^−/−^ mice demonstrated a greater than 5-fold increase in fecal oocyst shedding at 5dpi (Figure 5G). We note that parasite burden in *Irgm1/m3*^−/−^ mice was intermediate between WT and *Ifng*^−/−^ or *Stat1*^ΔIEC^ mice, which argues for additional IFN-γ/STAT1-mediated mechanisms for parasite control. However, these findings provide the first evidence for a role of IRGs in the IFN-γ-mediated mechanism of resistance to *Cryptosporidium*.

## DISCUSSION

The importance of IFN-γ in resistance to *Cryptosporidium* is well appreciated (Hayward et al., 2000; Leav et al., 2005; Pollok et al., 2001), but significant gaps exist in our understanding of the specific cells that produce and respond to this central cytokine in order to mediate parasite control. Previous studies have focused on NK cells as a potential innate source of IFN-γ in resistance to *C. parvum* (Barakat et al., 2009a; Rohlman et al., 1993); however, they predate the identification of other ILC populations. As confirmed in our studies, NK cells are largely present in secondary lymphoid tissues and the circulation, whereas the other ILC subsets are predominately tissue resident (Kim et al., 2016). Although there is marked phenotypic and functional plasticity for ILCs in the intestine (Gury-BenAri et al., 2016), our studies with *C. tyzzeri* suggested that ILC1s were the major early source of IFN-γ in the small intestine, although NK cells likely also contributed. In other models of intracellular infection, NK and ILC1-mediated resistance is transient, associated with the acute phase of infection but not sufficient for long term control (Park et al., 2019; Weizman et al., 2017). Thus, it was unexpected that although innate production of IFN-γ was not sufficient for parasite clearance in *Rag*^−/−^ mice, it did provide a significant level of long-term control of *C. tyzzeri*. Because of the lack of effective therapies to treat chronic cryptosporidiosis in patients with primary or acquired defects in T cell function (Flanigan et al., 1992; Navin et al., 1999; O’Hara et al., 2007), this model provides an opportunity to understand how ILC responses are maintained and whether they can be enhanced to mediate parasite clearance.

Previous studies have demonstrated that IL-12 and IL-18 are important in resistance to *C. parvum* (Bedi et al., 2015; Ehigiator et al., 2007; McDonald et al., 2006; Tessema et al., 2009; Urban et al., 1996). While DCs are a major source of IL-12 (Martinez-Lopez et al., 2015; Mashayekhi et al., 2011; Potiron et al., 2019), a recent study reported that, *in vitro*, *C. parvum* can directly activate the inflammasome and downstream caspase-1 in DCs, resulting in IL-18 secretion (McNair et al., 2018). In addition, IL-18 has been proposed to promote parasite control independent of IFN-γ in enterocyte cell lines (McDonald et al., 2006). In contrast, we found that enterocyte-intrinsic NLRP6 inflammasome activation was required for IL-18 mediated resistance to *C. tyzzeri in vivo* (Sateriale et al., 2021). Consistent with that observation, the studies presented here highlight enterocytes as a critical source of IL-18 which synergized with IL-12 to stimulate ILC1s to produce early IFN-γ. During infection with *Citrobacter rodentium*, an extracellular bacteria that attaches to the luminal surface of enterocytes, NLRP3 inflammasome activation leads to enterocyte-derived IL-18, which is required for host resistance (Liu et al., 2012; Munoz et al., 2015; Navabi et al., 2017). Thus, different sensors in enterocytes allow these cells to respond to diverse pathogens but converge on the processing of IL-18. In contrast, although *Salmonella* infects enterocytes, enteric neurons (not enterocytes) are the relevant source of IL-18 required for protection (Jarret et al., 2020). Intriguingly, patients with Hirschsprung disease have regions that lack distal bowel ganglia and are susceptible to *Cryptosporidium* (Sellers et al., 2018; Teitelbaum et al., 1989), but additional experiments will be required to determine whether the enteric nervous system is also a relevant source of IL-18 required for resistance to *Cryptosporidium*.

IFN-γ is a cytokine with wide ranging effects relevant to *Cryptosporidium* that include its ability to enhance antigen presentation and activation of anti-microbial activities within non-hematopoietic cells. Here, deletion of STAT1 was utilized to target the downstream effects of IFN-γ during *C. tyzzeri* infection. It is important to note that STAT1 is utilized by other IFNs and cytokines and there are reports that endogenous type I IFNs (Barakat et al., 2009b) and IFN-λ contribute to early resistance to *C. parvum* (Ferguson et al., 2019). Nevertheless, in the experiments performed with *C. tyzzeri*, mice deficient in STAT1 or IFN-γ showed similar susceptibility. Furthermore, the genes most strongly enriched in enterocytes from infected mice are more closely associated with IFN-γ, compared to IFN-α/ß or IFN-λ (Tretina et al., 2019). The finding that the loss of STAT1 in macrophage and DC populations did not impact susceptibility indicated that these populations were not important effectors of IFN-γ-mediated parasite control. In contrast, the inducible deletion of STAT1 in enterocytes established a cell-intrinsic role for STAT1 in resistance to *C. tyzzeri*. This pathway contrasts with *Salmonella*, where IFN-γ is required to restrict bacterial growth in macrophages but not enterocytes (Monack et al., 2004; Songhet et al., 2011).

Studies using human intestinal cell lines concluded that IFN-γ can inhibit growth of *Cryptosporidium*, but inhibitory effects *in vitro* are modest and the mechanisms that underlie parasite restriction are not understood (Khalil et al., 2018; Pollok et al., 2001). The identification of a role for Irgm1 suggests a potential overlap with mechanisms used to control other intracellular pathogens. Likewise, the upregulation of *Irgm1, Igtp* and *Iigp1* in enterocytes from infected mice highlights anti-microbial effectors that together with the GBPs can intersect with autophagy pathways involved in pathogen restriction (Coers et al., 2018). Although the loss of the GBPs on chromosome 3 (GBP 1, GBP2, GBP3, GBP5, and GBP7) had no detectable impact on host susceptibility, it does not rule out possible roles for the other GBPs located on chromosome 5. Of these, the expression of GBP6 transcripts was significantly enriched in epithelial cells from infected mice. Another possibility is that, although GBPs interact with vacuoles that contain other intracellular pathogens, perhaps the extra-cytoplasmic location of *Cryptosporidium* and the partitioning with a thick actin pedestal precludes this interaction, which would render the canonical GBP-dependent mechanisms of protection ineffective. The recent advances using enteroid-derived systems to culture *Cryptosporidium* (Heo et al., 2018; Wilke et al., 2019) should be useful to dissect how enterocytes utilize interferon-stimulated genes to directly clear or restrict growth of *Cryptosporidium* species.

The intestinal epithelium is a critical barrier where the immune system interacts with diverse microbial communities in the gut. Enterocyte interactions with the microbiome are central to homeostasis and the etiology of a variety of allergies and inflammatory conditions (Dahan et al., 2007; Eberl and Lochner, 2009; Peterson and Artis, 2014). There is an increased appreciation that different types of epithelial cells provide signals that help to coordinate the enteric nervous system, gut physiology and mucosal immunity in order to maintain tolerance. For extracellular pathogens, enterocytes promote clearance of the helminth *Trichuris muris* (Zaph et al., 2007) and the bacteria *Clostridium difficile* (Mamareli et al., 2019) and *Citrobacter rodentium* (Navabi et al., 2017). Less is known about how enterocytes respond to intracellular infections, although inflammasome-mediated extrusion of *Salmonella*-infected enterocytes limits bacterial spread (Rauch et al., 2017). Indeed, expulsion of infected enterocytes represents a conserved mechanism of pathogen resistance that is present in insects and higher vertebrates (Ayyaz and Jasper, 2013; Chatterjee and Ip, 2009; Lee et al., 2016). The studies presented here identified enterocytes as a critical source of IL-18 required for ILC activation and as key drivers of IFN-γ mediated resistance to an important enteric pathogen. Thus, *C. tyzzeri* is a valuable natural model of enteric infection that affects intestinal physiology and nutritional status which can provide novel insights into the role of enterocytes in recognition of and resistance to infection.

## SUPPLEMENTAL FIGURES

**Supplemental Figure 1.**
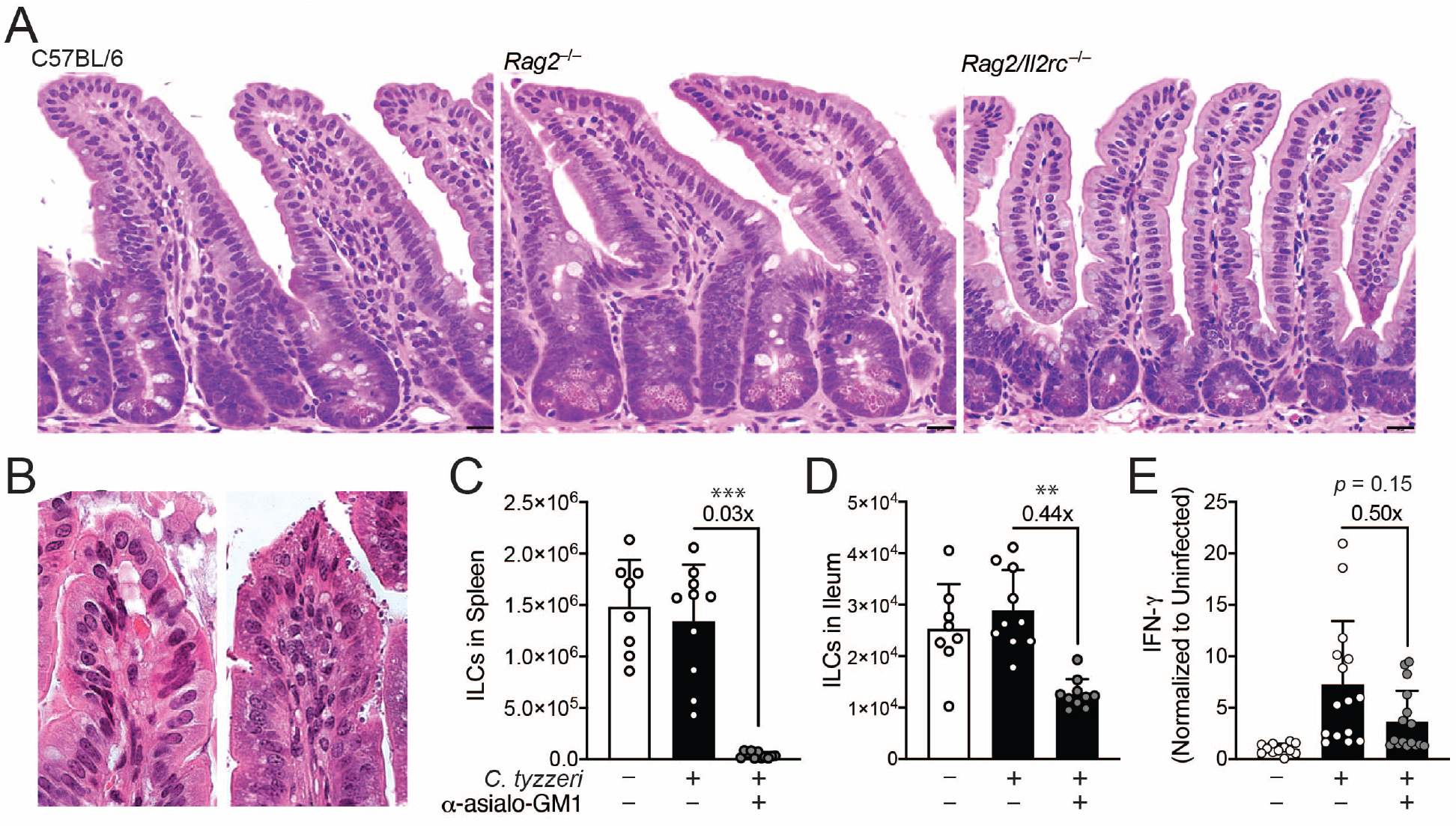
(Refers to Figures 1 and 2) (A) Representative H&E staining of ileal sections from naïve C57BL/6 (left), *Rag2*^−/−^ (center), and *Rag2*^−/−^*Il2rc*^−/−^ mice. (B) Representative H&E of an illeal villus from Rag2-/- mice treated with IgG (left) or α-IFN-γ (right). (C-E) *Rag2*^−/−^ were treated with α-asioloGM1 (see Methods). Total numbers of ILCs (CD45.2^+^CD3- NKp46^+^NK1.1^+^) from the spleen (C) or ileum (D) are shown in cumulative bar graphs. IFN-γ was measured from the supernatants of ileal biopsies by ELISA (see Methods) and are shown in (E).

**Supplemental Figure 2.**
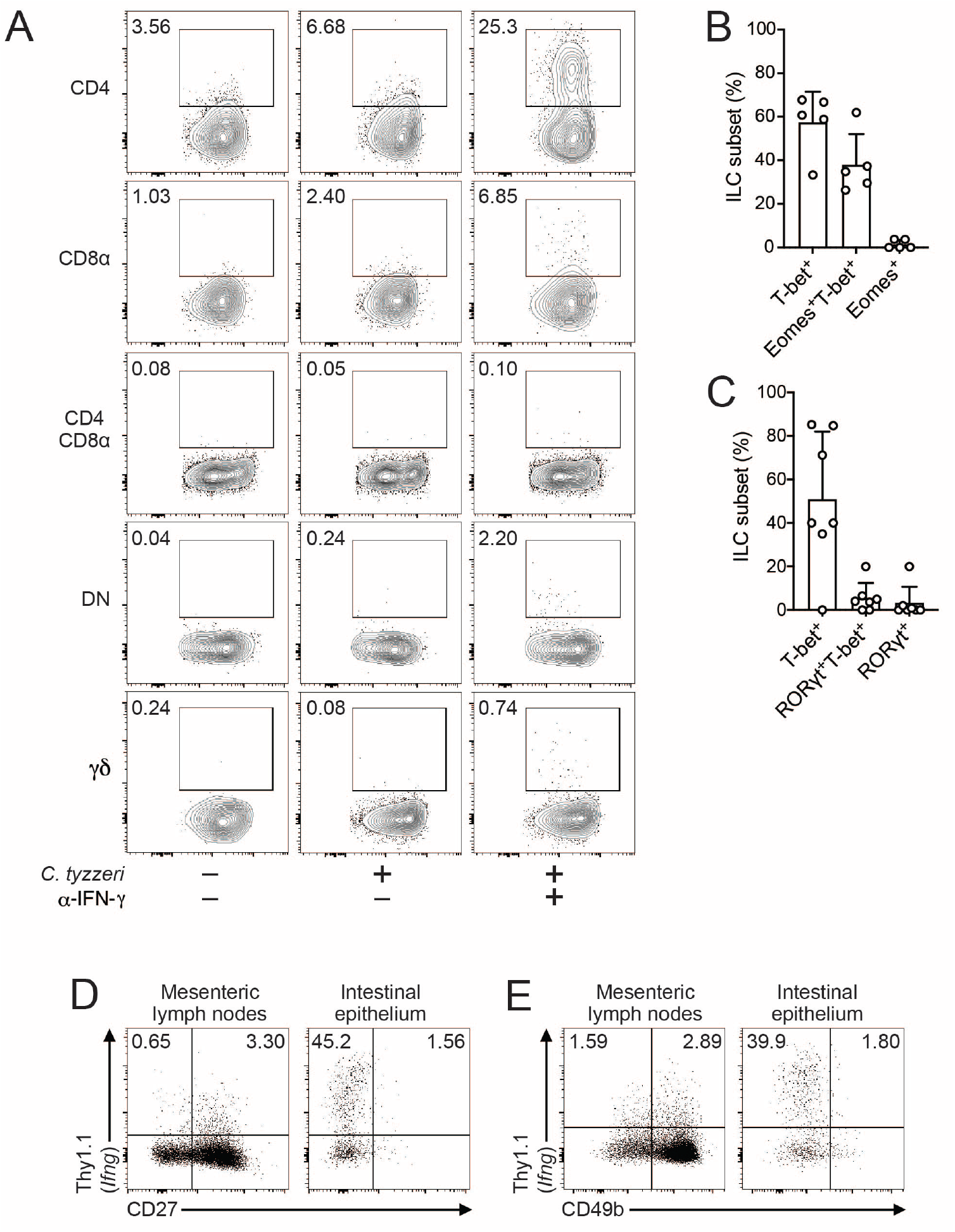
(Refers to Figure 3) (A) Representative flow plots of Thy1.1 expression in intestinal T cell subsets from Ifng-Thy1.1 KI mice. Left: uninfected, center: 4dpi with *C. tyzzeri*, right: 4dpi with *C. tyzerri* and treated with α-IFN-γ. (B-C) Summary bar graphs of % of ileal ILC subsets from naive mice. (D-E) Representative flow plots of ILCs 4dpi (gating same as Figure 3).

**Supplemental Figure 3.**
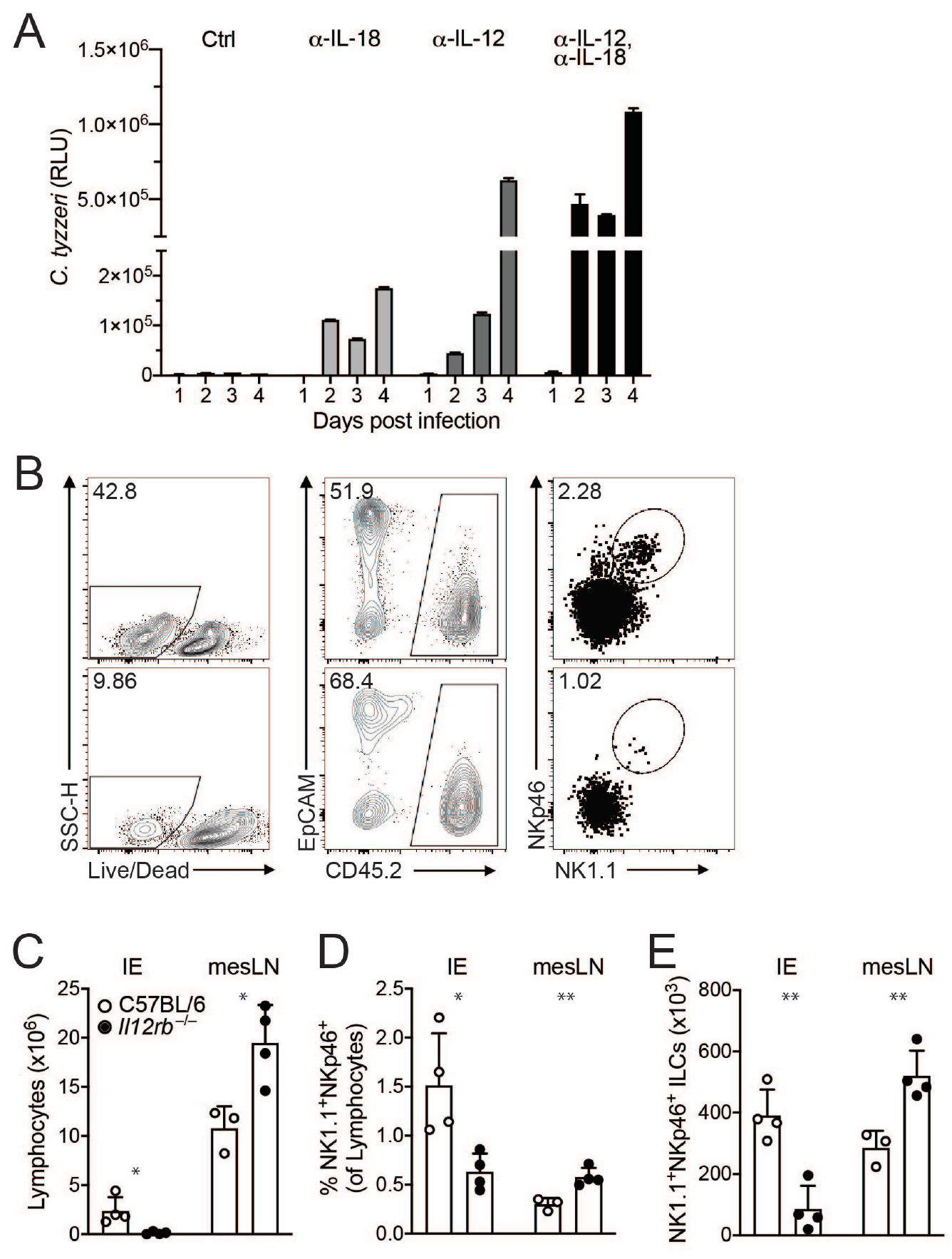
(Refers to Figure 4) (A) C57BL/6 mice treated with α-IL-18, αIL-12, or both were infected with *C. tyzzeri* and oocyst shedding in feces was monitored. Representative of 3 experimental replicates. (B) Representative flow data of cells from the ileum of C57BL/6 or *Il12rb*^−/−^ mice, gated on live cells (left), CD45.2^+^ (center), NK1.1^+^NKp46^+^ (right). (C-E) Cumulative bar graphs of total lymphocytes (C), frequency of NK1.1^+^NKp46^+^ (D) and total ILC numbers (E).

**Supplemental Figure 4.**
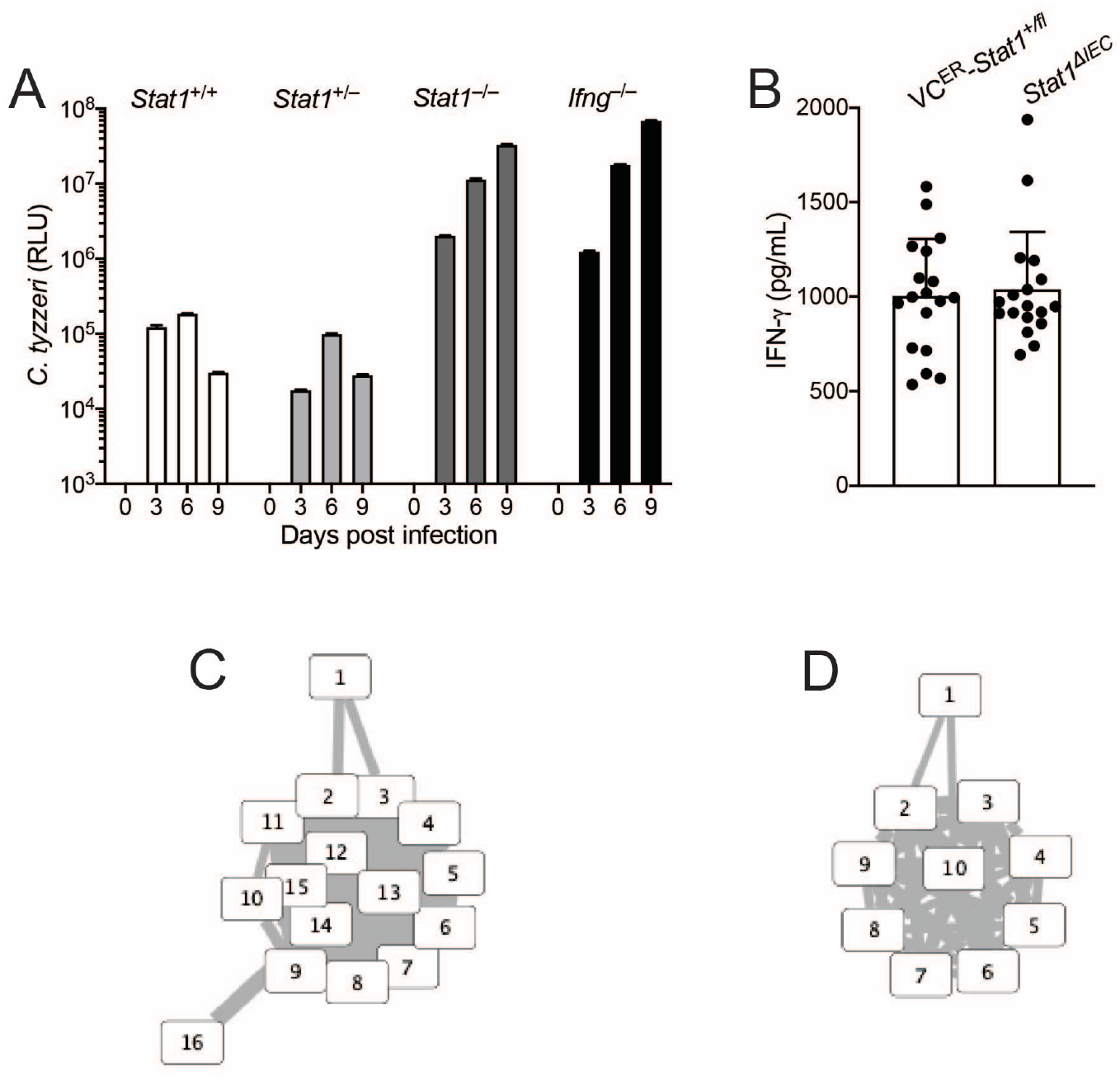
(Refers to Figure 5) (A) *Stat1*^−/−^ mice, littermate controls (*Stat1*^+/+^ and *Stat1*^+/-^) and *Ifng*^−/−^ mice were infected with *C. tyzzeri* and oocyst shedding was monitored in feces. (B) IFN-γ was measured from the supernatants of ileal biopsies by ELISA. (C-D) GSEA clusters of mitochondrial (C) and ribosomal/protein translation pathways (D).

## Notes

This work was supported in part by funding from the Bill and Melinda Gates Foundation (OPP1183177) to BS and through grants from the U.S. National Institutes of Health to BS & CAH (R01AI148249), R21 AI139895 to CAH, and fellowships and career awards to JG (T32AI055400), AS (K99AI137442), and AG (T32A1055400) and a Swiss National Science Foundation fellowship (191774) to SB.

### Competing Interest Statement

The authors have declared no competing interest.

